# High transcriptional error rates vary as a function of gene expression level

**DOI:** 10.1101/554329

**Authors:** K.M. Meer, P.G. Nelson, K. Xiong, J. Masel

## Abstract

Errors in gene transcription can be costly, and organisms have evolved to prevent their occurrence or mitigate their costs. The simplest interpretation of the drift barrier hypothesis suggests that species with larger population sizes would have lower transcriptional error rates. However, *Escherichia coli* seems to have a higher transcriptional error rate than species with lower effective population sizes, e.g. *Saccharomyces cerevisiae*. This could be explained if selection in *E. coli* were strong enough to maintain adaptations that mitigate the consequences of transcriptional errors through robustness, on a gene by gene basis, obviating the need for low transcriptional error rates and associated costs of global proofreading. Here we note that if selection is powerful enough to evolve local robustness, selection should also be powerful enough to locally reduce error rates. We therefore predict that transcriptional error rates will be lower in highly abundant proteins on which selection is strongest. However, we only expect this result when error rates are high enough to significantly impact fitness. As expected, we find such a relationship between expression and transcriptional error rate for non C➔U errors in *E. coli* (especially G➔A), but not in *S. cerevisiae*. We do not find this pattern for C➔U changes in *E. coli*, presumably because most deamination events occurred during sample preparation, but do for C➔U changes in *S. cerevisiae*, supporting the interpretation that C➔U error rates estimated with an improved protocol, and which occur at rates comparable to *E. coli* non C➔U errors, are biological.

## Main Text

Errors are costly, and we therefore expect natural selection to reduce their rate. However, selection cannot achieve everything. In particular, it is only able to purge deleterious mutations when their selection coefficient *s* is significantly greater than one divided by the “effective population size”. This numerical limit to selection may reflect not just the number of individuals in a population, but also competing selection at linked sites (Good and Desai 2014; Lynch 2007). The “nearly neutral theory” holds that deleterious mutations close to this limit are abundant (Ohta 1973), and the “drift barrier hypothesis” holds that differences in the precise location of this limit explain important differences among species (Lynch 2007). For example, codon usage bias is stronger in species believed to have higher effective population sizes (Vicario et al. 2007), indicating stronger selection to purge slightly deleterious synonymous mutations.

Rajon and Masel (2011) highlighted the distinction between a “global” solution that ameliorates a problem at many loci at once, and a set of “local” solutions that solve them one at a time. Because mutations affecting single loci are likely to have smaller fitness consequences than mutations with genome-wide effects, the drift barrier forms a more formidable barrier to local solutions than it does to global solutions. When local solutions evolve (in populations with large effective population sizes), they can obviate the need for global solutions. This yields the counterintuitive prediction that when global solutions are examined, it may be species with low effective population sizes that show the most extreme adaptations. Specifically, rates of error in transcription and translation could be higher in species with high effective population sizes, since reducing error rates by kinetic proofreading is a costly global solution (Rajon and Masel 2011).

Here we focus on mistranscription errors, where during transcription, the wrong nucleic acid is incorporated at a single site. This can lead to non-functional proteins, incurring three types of costs. First is the energetic cost of futile transcription and translation (Wagner 2007); which can be significant in bacteria with large population sizes (Lynch and Marinov 2015; Petrov and Hartl 2000). Second, there is the opportunity cost of not using ribosomes to make other gene products (Dekel and Alon 2005; Kafri et al. 2016; Scott et al. 2014). Third, there is the cost of disposing of a misfolded and potentially toxic protein (Drummond and Wilke 2009; Geiler-Samerotte et al. 2011; Tomala and Korona 2013). Rajon and Masel (2011) predicted that in populations with smaller effective population sizes and more loci, costly proofreading might evolve to reduce the rate of mistranscription and hence the frequency with which these three costs are born, while in populations with very large effective population sizes and fewer loci, local solutions might evolve to reduce the cost of each mistranscription event, allowing their rate to stay high.

This prediction seems to have been confirmed for mistranscription (Xiong et al. 2017), whose rate of 8.2×10^−5^ in *Escherichia coli* (Traverse and Ochman 2016b) is far higher than that in *Saccharomyces cerevisiae* (3.9×10^−6^) (Gout et al. 2017) or *Caenorhabditis elegans* (4.1×10^−6^) (Gout et al. 2013), which have lower effective population sizes. Indeed, the rate is higher even than that of *Buchnera aphidicola* (4.7×10^−5^) (Traverse and Ochman 2016b). *Buchnera* is a highly mutationally degraded species in which the drift barrier is an obstacle to the maintenance of fidelity in many other important cellular functions (McCutcheon and Moran 2012); this high rate in *Buchnera* may thus indicate that the drift barrier forms an obstacle even to global solutions (Xiong et al. 2017). All these error rates except for that of *C. elegans* (Gout et al. 2013), were estimated using Cir-Seq (Acevedo and Andino 2014), and should therefore be comparable, although sample preparation techniques differ in vulnerability to deamination.

If the drift barrier theory of Rajon and Masel (2011) explains the high rate of mistranscription in *E. coli*, this implies that selection in *E. coli* must be potent enough to be sensitive to the consequences of transcription errors in a local (i.e. site-specific) way, not just to its global rate. Local solutions to mistranscription fall into two categories: local robustness to the consequences of mistranscription when it occurs (this evolved robustness is hypothesized to be responsible for permitting globally high mistranscription rates), and locally reduced mistranscription rates at the sites most sensitive to it.

Here we test whether selection is able to maintain locally lower transcriptional error rates in highly expressed genes. Selection to purge deleterious mutations is generally more effective in highly expressed genes, as evidenced, for example, by stronger codon bias (Cutter and Charlesworth 2006; Duret and Mouchiroud 1999; Ran et al. 2014; Sharp et al. 2010), which lowers translational error rates (Zhang et al. 2016). Somatic mutations (Frigola et al. 2017), alternative transcriptional start sites (Xu et al. 2019), post-transcriptional modifications (Liu and Zhang 2018a, b), alternative mRNA polyadenylation (Xu and Zhang 2018), and translation errors (Mordret et al. 2019) also occur at lower rates at sites where they are likely to have larger effects. We similarly predict that because high mistranscription rates matter more for highly expressed genes, highly expressed genes should evolve a lower rate of mistranscription. We make this prediction for *E. coli*, where mistranscription rates are globally high and thus so is local selection pressure. In contrast, we do not expect a relationship between expression level and mistranscription rate in *S. cerevisiae*, where mistranscription rates are globally much lower.

Mistranscription rate data in *E. coli* were taken from Traverse and Ochman (2016a), who used Cir-Seq (Acevedo and Andino 2014) to distinguish mistranscription events from sequencing errors. Within the largest and highest-quality batch of their data (see Methods), data from four experimental conditions (minimal vs. rich media, and midlog vs. stationary phase) were sometimes analyzed separately and sometimes pooled. Mistranscription rates are much higher for C➔U substitutions: ~10^−4^ rather than ~10^−5^ for other mistranscription types. Since C➔U changes are more sensitive to preparation artifacts (Chen et al. 2014), i.e. they may not be mistranscription errors, we excluded them from most of our analysis.

To further ensure the data quality, we exclude “hotspot” nucleotide sites experiencing significantly (*p*<10^−9^) more errors of one type than expected from our model fitted as described below. This eliminates recent mutations, inaccurate mapping of reads to the genome, or other artifacts of the experiment or pipeline, as well as any sites subject to programmed post-transcriptional RNA editing. We excluded 5 protein-coding and 2,390 non-coding sites that met this “hotspot” criteria for at least one experimental condition. The high rate of apparent mistranscription hotspots in non-coding genes has been interpreted (Traverse and Ochman 2016a) as a consequence of *E. coli* having multiple polymorphic rRNA operons, making mapping of reads inaccurate. We therefore restrict our analysis to protein-coding genes.

We modeled the number of errors observed per nucleotide site as count data, using a generalized linear model. The number of errors expected is the product of the number of observations of that nucleotide site, and the modeled mistranscription rate, the latter a linear function of log protein abundance, experimental condition, and substitution type (see Methods). The dependence on protein abundance (Figure 1; slope of 0 rejected from Eq. 1 model with *p*=2×10^−14^) supports our prediction from drift barrier theory, a result that gets slightly stronger if we omit our hotspot removal procedure. The 11 non-C➔U substitution types have substantially different mistranscription rates (Supplementary Figure S1); fitting different intercepts for each type (while leaving their slopes the same) is strongly supported for inclusion in our Eq. 1 model (*p* = 2×10^−16^).

**Figure 1.**
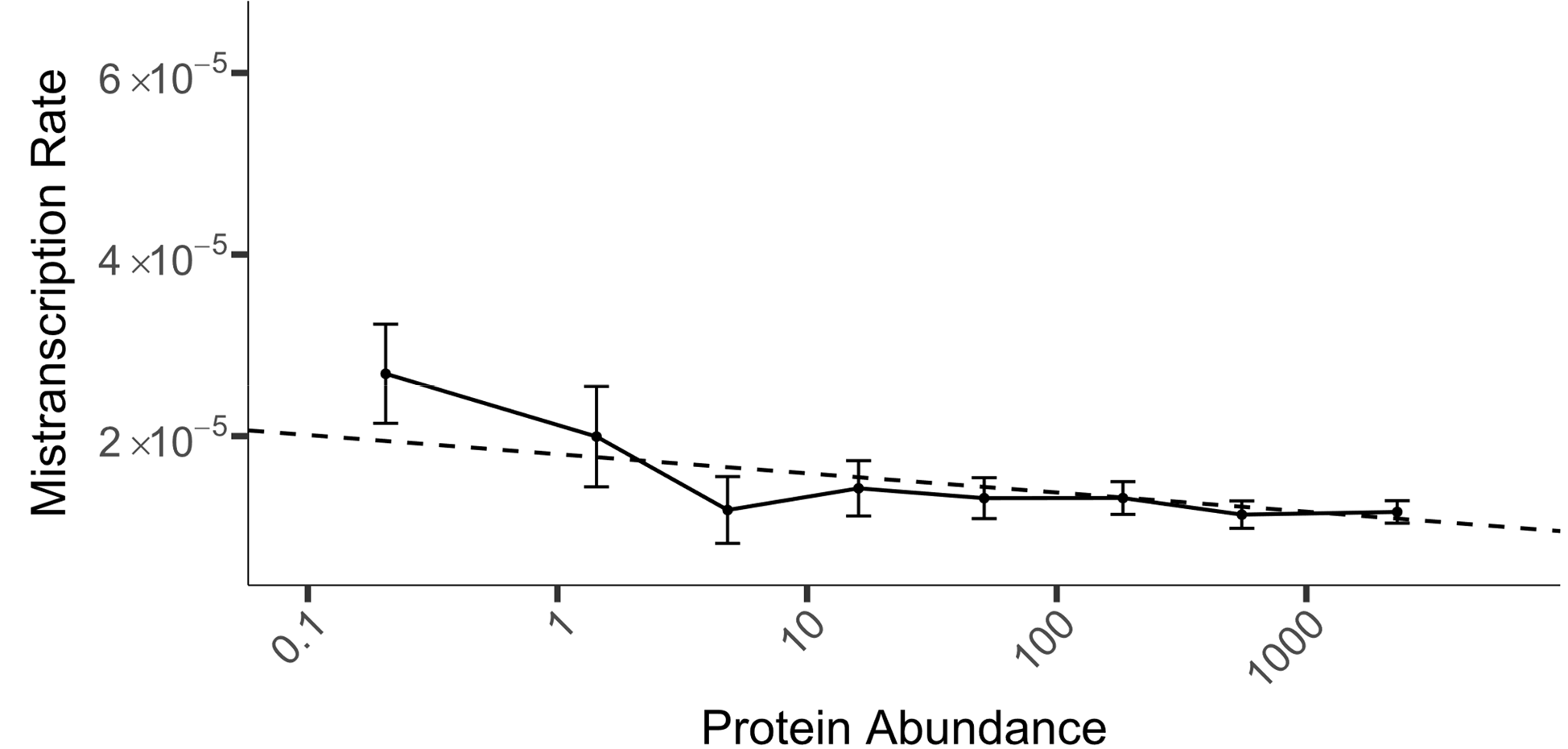
Highly expressed *E. coli* genes are subject to lower mistranscription rates. The dashed line shows the Eq. 1 model applied to the 11 non C➔U substitution types, in which both condition and substitution type affect the intercept but not the slope, plotted as a weighted average over conditions and substitutions, with weights proportional to the frequencies of opportunity to occur (i.e. by the numbers of reads of sites with A/C/G/U). Solid line show the pooled data, binned by protein abundance as described in the Methods, and plotted according to mean protein abundance and the mean and 95% CI of the mistranscription rate within each bin. Data were divided into 10 bins; because of the limited availability of reads for low-expression genes, data within the first three bins were pooled. Note that mistranscription rate is per possible error, so the total mistranscription rate per nucleotide is around three times larger.

Different intercepts for different experimental conditions are also supported, in addition (*p* = 1.5×10^−3^). Fitting different slopes for each experimental condition only marginally improves the fit relative to our Eq. 1 model (*p* = 0.052), mostly attributable to a steeper slope in the minimal-static condition, which had far fewer data points than the other conditions (Figure S2).

Standing out from results on all non C➔U error types in Figure S1, and shown in Figure 2, is the fact that G➔A errors depend more strongly on protein abundance than other error types do (*p*=3×10^−4^, Eq. 2 as improvement on Eq. 1). A separate model fit to G➔A error data only, gives a slope of −2.9×10^−6^ (95% CI of −1.6×10^−6^ to −4.2×10^−6^) with log_10_ protein abundance, i.e. there are 1.6 to 4.2 fewer G➔A errors per million G transcription events per 10-fold increase in expression, against a background of about 20-40 errors per million G transcription events. To ensure that the non-zero slope of Figure 1 is not driven solely by G➔A errors, we repeated the analysis for the 10 error types, i.e. excluding both C➔U and G➔A (Figure 2, right). This yields a slope of −8.4×10^−7^ (*p*=1×10^−7^) with log_10_ protein abundance, with a 95% confidence interval corresponding to 0.4 and 1.2 fewer expression errors per million opportunities per 10-fold increase in expression.

**Figure 2.**
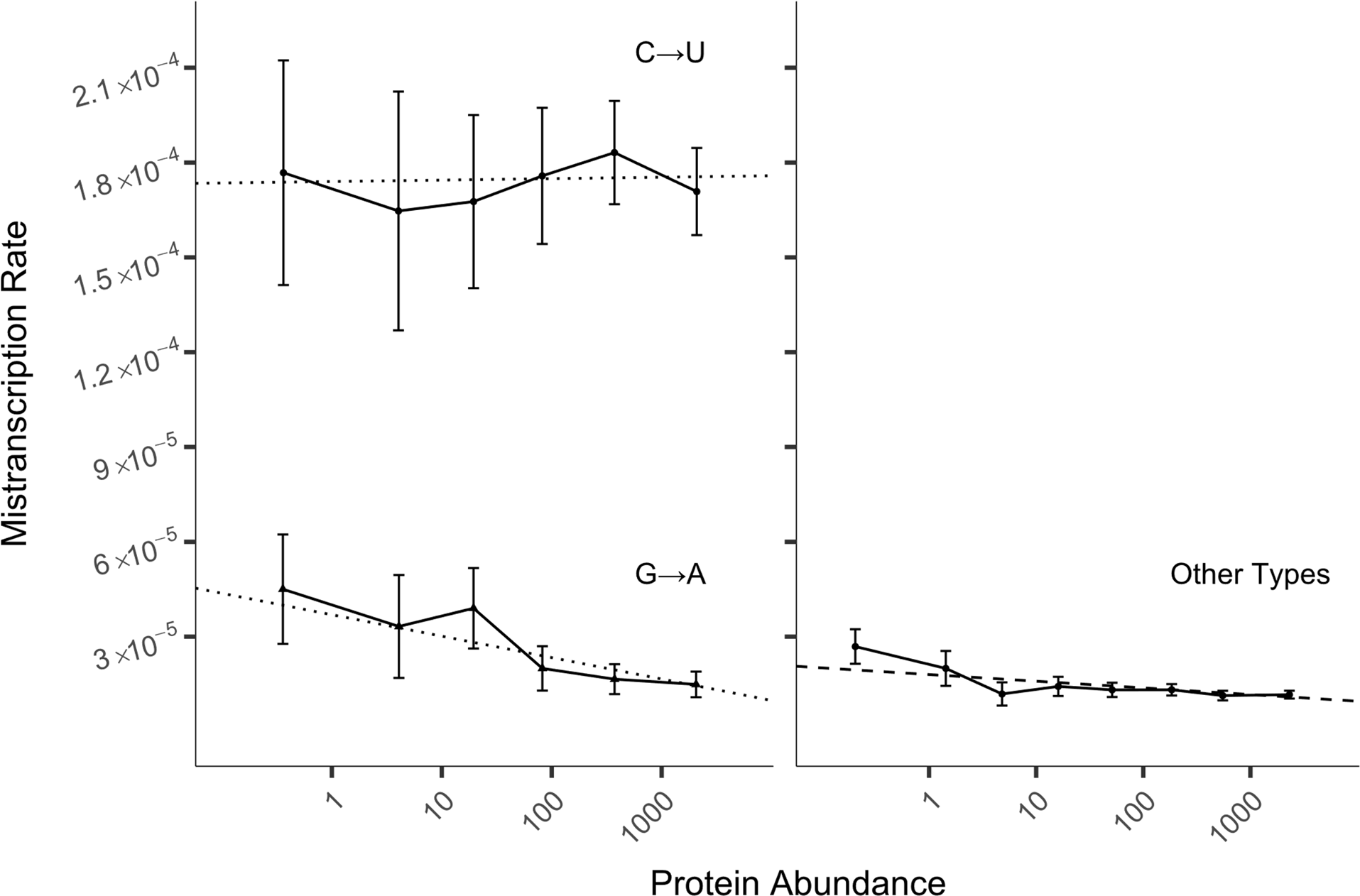
C➔U errors in *E. coli* are mostly artifacts, G➔A depend most strongly on protein abundance, but the other 10 error types also show dependence. Dotted lines (left) show linear models with both the slope and intercept fitted separately for each error type using data pooled across all four conditions; for a comparison of all 12 error types, see supplemental Figure S1. The C➔U slope is not different than 0 (*p*=0.91). The dashed line (right) shows an Eq. 1 model in which the slope is the same across all 10 error types (non-C➔U, non-G➔A). To display this model, we averaged the intercept over the four conditions, weighted according to the numbers of reads in each condition. Solid lines show the mean mistranscription rates, binned by protein abundance as described in the Methods, plotted according to mean protein abundance within each bin; error bars show 95% CI. Data were divided into 8 bins; because of the limited availability of reads for low-expression genes, data within the first three bins were pooled. Note that mistranscription rate is per possible error, so total mistranscription rate per nucleotide is around three times larger.

Traverse and Ochman (2016a) reported that mistranscription errors were more commonly synonymous (32%) than would be predicted if errors occurred at random across the genome (24%). When controlling for the effects of substitution type, condition, and protein abundance in our Eq. 2 model of mistranscription rates, the synonymous vs. non-synonymous status of the potential mistranscription error did not predict the error rate (*p* = 0.89). Indeed, following our data processing and quality filters, the overall frequency with which a mistranscription error was synonymous was 23.4%, suggesting that the previously reported excess of synonymous mistranscription events was due to data quality issues. In any case, whatever molecular mechanism is responsible for variation in mistranscription rates, it seems to act at the level of the gene rather than at the level of the nucleotide site.

Molecular chaperones play a critical role in mitigating the harm from mistranscription by reducing misfolding. Genes that are chaperone clients might tolerate higher mistranscription rates. Alternatively, sensitivity to mistranscription might select both for a lower mistranscription rate and chaperone use. We found no support for either hypothesis; adding an intercept term for *GroEL* chaperonin use was not a significant improvement on top of our Eq. 2 model (*p* = 0.085). We also tested other predictors including gene length, absolute position of a locus (number of nucleotides from the start of gene), and relative position of a locus (absolute position / total gene length), but neither slope nor intercept were significantly different from 0 (i.e. *p*>0.05) for any of the three metrics.

As discussed in the Introduction, Cir-Seq data on the yeast *S. cerevisiae* indicates a much lower mistranscription rate than *E. coli* (Gout et al. 2017), suggesting that it uses a global solution, reducing site-specific selection pressures on mistranscription rates. We therefore do not predict a relationship between gene expression and local mistranscription rate in this species, and do not find one for the 11 non C➔U substitution types (Figure 3 bottom; *p*=0.2 in our Eq. 1 model controlling for substitution type as a fixed effect).

**Figure 3.**
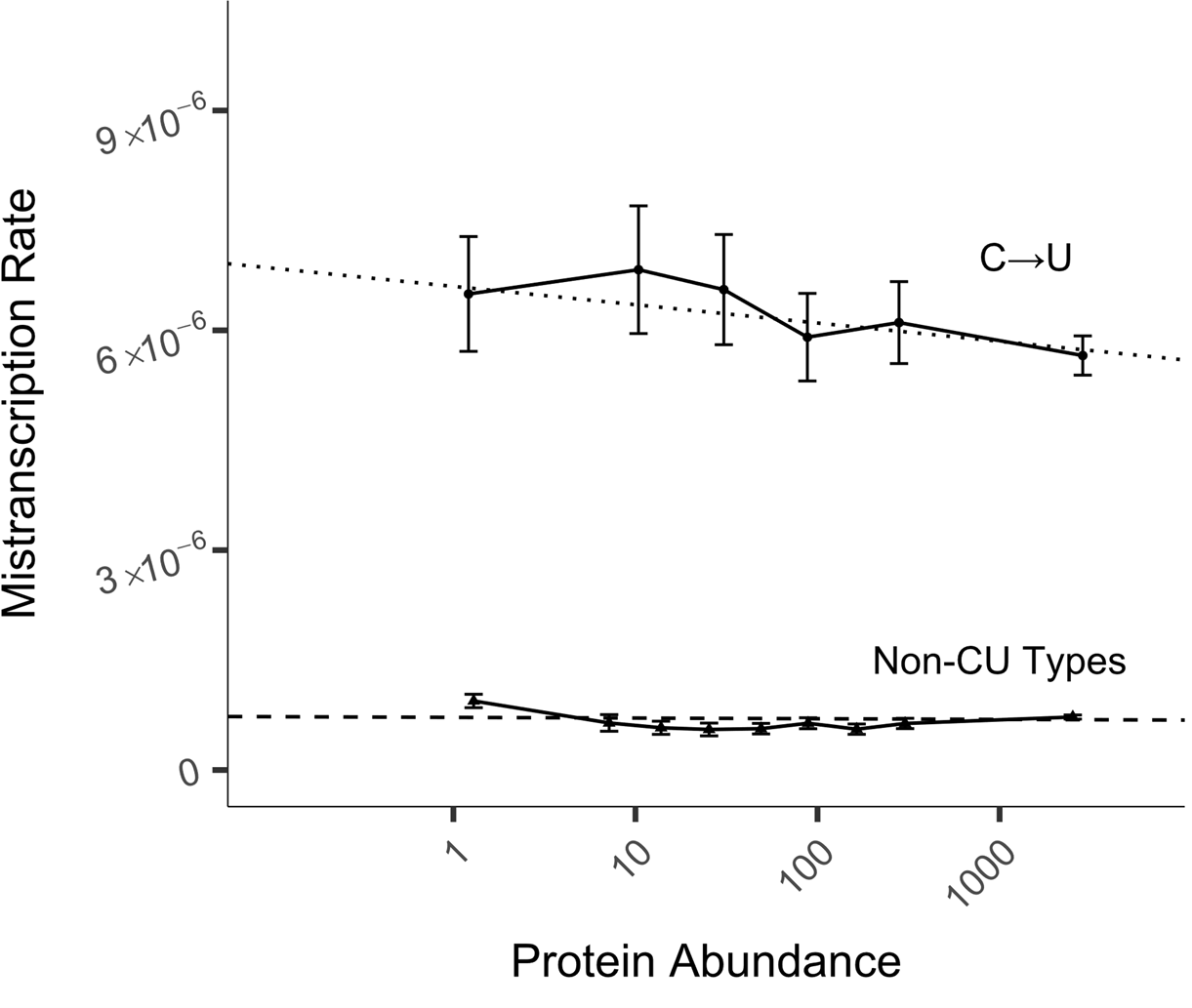
In *S. cerevisiae*, only C➔U mistranscription errors depend on protein abundance. Dashed line shows a linear model fitted to a pooled dataset of the 11 non-C➔U substitution types. Dotted line shows a linear model fitted to C➔U data alone. To display non-C➔U model fit, we took a weighted average of the intercept over substitution types as a function of the frequencies of opportunity to occur. Solid line shows pooled data, binned by protein abundance as described in the Methods, and plotted according to mean protein abundance and the mean and 95% CI of the mistranscription rate within each bin. C➔U data were divided into 8 bins; because of the limited availability of reads for low-expression genes, data within the first three bins were pooled. For non-C➔U data, two out of 10 bins were pooled. Note that mistranscription rate is per possible error, so total mistranscription rate per nucleotide is around three times larger. All 12 error types are shown separately in Supplementary Fig. S3.

However, C➔U substitutions, which occur at much higher rates than other substitution types and hence are subject to more selection even in *S. cerevisiae*, are less frequent for highly abundant proteins (Fig. 3 top; *p*= 0.006 for non-zero slope on a C➔U equivalent of Eq. 2). This confirms that the protocol of Gout et al. (2017) succeeded in avoiding deamination events during sample preparation (which should not depend on protein abundance), where that of Traverse & Ochman (2016a) did not.

The high rate of mistranscription errors in *E. coli* came as a surprise to many (Traverse and Ochman 2016a, b). This naturally raises the hypothesis that it is the data that are in error. While the Cir-Seq technique is effective in preventing sequencing errors from inflating estimated mistranscription rates (Acevedo and Andino 2014), it does not eliminate artifacts of the sample preparation and analysis such as mutations occurring during the Cir-Seq experiment, nor inaccurate mapping of reads to the genome. While these could artificially inflate estimated mistranscription rates, we are not aware of any plausible mechanism by which the degree of such inflation would be a function of protein abundance. Our results thus confirm the credibility of the data, and hence of the statement that *E. coli* has a strikingly high non-C➔U mistranscription rate. After applying our quality filters, we calculate the total rate of all non-C➔U errors as 4.1×10^−5^ per site, or 8.6×10^−5^ if C➔U errors are also included. In contrast, in *S. cerevisiae*, we calculate from the data of Gout et al. (2017) a non-C➔U mistranscription rate of 2.3×10^−6^, or 3.5×10^−6^ with the C➔U error type included.

The dependence of the *E. coli* mistranscription rate on the strength of selection (as reflected by protein abundance), but not the *S. cerevisiae* mistranscription rate, is consistent with proposed drift barrier explanations (McCandlish and Plotkin 2016; Rajon and Masel 2011; Xiong et al. 2017). In particular, *E. coli* is smaller and is generally accepted to have a larger effective population size than *S. cerevisiae*. *E. coli* also has fewer loci, occurring within 4453 genes in K-12 (Riley et al. 2006) compared to 5178 genes in *S. cerevisiae* (Engel et al. 2014), which makes it easier to evolve robustness at each one. What is more, the average *E. coli* mRNA produces about 540 proteins out of a total of 2.5 × 10^6^ per cell (Lu et al. 2007), i.e. 0.02% of the proteome, which is twice as much as the average yeast mRNA producing 5600 proteins out of a total of 5 × 10^7^ per cell (Lu et al. 2007), i.e. 0.01% of the proteome. While a typical yeast mRNA has a longer half-life and so makes proteins over a longer time (6.7 vs. 27.4 minutes; Siwiak and Zielenkiewicz 2013), the magnitude of this should not be enough to counteract all other factors making local solutions easier to evolve in *E. coli*.

We have shown that local mistranscription rates vary in a systematic way on a per-gene basis, but have not determined the mechanisms by which expression error rates vary. Mistranscription rates are affected by local sequence characteristics such as long mononucleotide repeats (Ackermann and Chao 2006; Gu et al. 2010) and at the gene level by the presence or absence of specific RNA polymerase subunits (Thomas et al. 1998; Walmacq et al. 2009) or transcription factors (Bubunenko et al. 2017; Irvin et al. 2014; Roghanian et al. 2015). Our finding that G➔A errors depend more strongly on expression than do other error types in E. coli suggests that GreA, which specifically reduces G➔A transcription errors (Traverse and Ochman 2018), may be a likely mechanistic candidate.

We have also shown that the local mistranscription rates even of highly expressed *E. coli* genes are higher than the global mistranscription rate in *S. cerevisiae*, suggesting that *E. coli* genes are somehow more robust to the consequences of mistranscription than are *S. cerevisiae* genes. However, the robustness associated with *E. coli*’s global solution is not so complete as to eliminate selection for locally lower mistranscription rates in the genes subject to the strongest selection, leading to the trend detected here.

## Methods

Scripts used in these analyses are available at https://github.com/MaselLab/Meer-et-al-Transcriptional-Error-Rates.

### E. coli mistranscription data

Pre-processed data were obtained from Traverse and Ochman (2016a), that included how many times each of the 4,641,652 nucleotide loci in the K-12 MG155 reference genome (GenBank accession: NC.000913.3) was observed, and how often each nucleotide was seen there. We assigned these loci to 4,140 protein coding genes and 178 non-coding genes using the annotation of GenBank accession NC.000913.3. We analyzed the 3,935,551 nucleotide loci within annotated non-overlapping protein-coding ORFs, and 47,344 nucleotide loci from non-coding genes based on annotated ‘start’ and ‘stop’ positions. We excluded any sites that were present in overlapping genes, as we could not assign a single error rate or protein abundance in such cases.

Traverse and Ochman (2016a) data were obtained in multiple batches (referred to as “replicates” in their data tables), with results reported only on two of the batches. Batch #2 had approximately half as much data and twice the error rate of batch #1, so we restrict our analysis to batch #1 only. Combining the data from each of the four experimental conditions (minimal vs. rich media, and midlog vs. stationary phase) within batch #1 effectively yielded 15,742,204 protein-coding sites and 189,376 non-coding sites, where “site” is used here as shorthand for condition×nucleotide locus, i.e. to describe the set of reads of a nucleotide locus within just one experimental condition.

We excluded any site that had no reads and any protein-coding transcript site with no protein abundance measure, leaving 5,994,463 coding and 182,233 non-coding sites. Each site can experience three different substitution error types (e.g. C➔U, C➔A, and C➔G), which we treated separately, yielding 17,983,389 coding and 546,699 non-coding “possible errors” for analysis. Note that data for the three alternative errors at the same site are not, strictly speaking, independent, because the occurrence of one error reduces the denominator for the other two. However, at low error rates, this effect is negligible.

Mutations occurring during the Cir-Seq experiment, inaccurate mapping of reads to the genome, or other artifacts of the experiment or pipeline can result in the appearance of mistranscription “hot spots” that are best removed. We calculated the likelihoods of seeing that many or more errors for each of the 18,530,088 possible errors being analyzed, using a significance cutoff of 10^−9^ to ensure that only 10^−9^×18,530,088=0.02 possible errors are falsely excluded, or potentially more if there is genuine biological variation in mistranscription rates beyond that captured by our linear model. We calculated likelihoods from a cumulative binomial distribution based on the number of reads at that site and the rate of error expected at that site from our model. When a possible error was excluded with likelihood < 10^−9^, we excluded the entire nucleotide locus (i.e. all three possible substitutions in all four conditions). We performed an iterative procedure, first fitting a model of constant error rate for all non C➔U errors and a separate error rate for C➔U errors, using expectations from this model to exclude outliers, then using the cleaned-up data to develop a more sophisticated error rate model of all conditions/substitution types, and using the revised expectations from this model to update which loci should be excluded etc. until convergence. In the final iteration, one or more possible errors was determined to be an outlier at 5 protein-coding and 2,390 non-coding loci. For protein-coding outliers, we excluded all possible errors at each of the 5 outlier loci, i.e. up to 60 possible errors (3 possible errors at 5 loci in 4 conditions). Some sites had no transcript reads in some conditions, resulting in only 48 rather than 60 possible errors being excluded by this procedure, leaving 17,983,341 possible errors in protein-coding transcript regions for analysis. Excluding C➔U substitutions, due to their significantly higher error rate and likelihood of occurring post-transcriptionally, further reduced this to 16,466,559 non-C➔U possible errors for analysis.

### S. cerevisiae mistranscription data

Similarly pre-processed transcript data were obtained from Gout et al. (2017), who recorded how many times each nucleotide locus was observed in the S288C reference genome (GenBank accession: GCA_000146045.2), to which the wild-type BY4741 strain used in their experiment is very closely related. Only one experimental condition was used in this study. Using the same methodology as for the *E. coli* data, we used the accession to assign nucleotide sites to the 5,983 protein-coding nuclear gene regions based on the annotated ‘start’ and ‘stop’ positions. This process identified 8,853,931 nucleotide loci within annotated protein-coding ORFs, resulting in 26,561,793 possible errors for analysis.

Excluding any transcript site without reads or with unreported or zero protein abundance left us with 18,649,818 possible errors. Using our outlier detection protocol, we identified 44 loci containing possible errors as outliers and excluded all possible errors at the associated loci (132 possible errors in total), leaving 18,649,686 possible errors for analysis.

C➔U errors were also identified as having a substantially higher error rate in the yeast data (1.8×10^−5^ versus 2.3×10^−6^ for other mistranscription types), and were excluded from some analyses, resulting in 17,394,875 non-C➔U possible errors.

### Protein abundance data

Integrated protein abundance data were taken from PaxDB (Wang et al. 2015).

### GroEL client status

We labelled the 1,929,741 possible errors associated with 252 *E. coli* proteins as having GroEL client status, based on the identification of those proteins by Kerner et al. (2005) as specific interactors with the GroEL chaperonin.

### Statistical model

We modeled the error rate at site *i* within gene *j* as a linear function of the log-abundance of protein *j*, i.e.

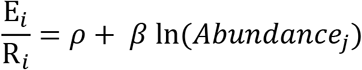

where E_*i*_ is the number of reads containing a particular error and R_*i*_ is the total number of reads at that nucleotide site.

To better model the error function in the linear model, we multiply both sides by R_*i*_:

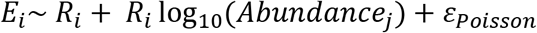

The observed number of errors E_*i*_ has the properties of count data, and so can be modeled as a sample from a Poisson distribution. We fitted the statistical model above using a generalized linear model function in R (glm, stats package), specifying the family of the model as “poisson(link = identity)”. For *E. coli*, experimental condition and type of error (excluding C➔U) were added as fixed effects to yield:

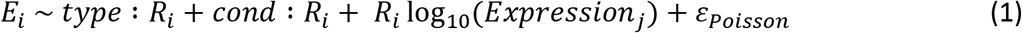

Slope as a function of expression level can also be made dependent on type and/or condition. For *S. cerevisiae*, the condition term does not apply, and expression was not supported as predictive in the model. For *E. coli*, a separate slope for G➔A errors was supported, yielding

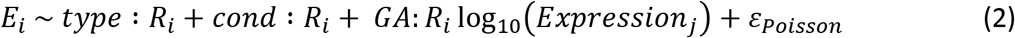

P-values associated with adding or removing terms to Eq. 1 or Eq. 2 models were obtained using the anova command with the Chisq option to compare nested models in R, as given throughout the text, sometimes manually correcting the number of degrees of freedom.

### Data Binning

We binned data by protein abundance for visualization and comparison to the fitted models. All possible errors were sorted by the abundance value of the corresponding protein. Bin boundaries were evenly spaced along our log-abundance axis between the 5% quantile and the 95% quantile, with data beyond these quantiles included in the edge bins. For each bin, one point was plotted with y-value equal to the mean and 95% confidence interval of the mistranscription rate and an x-value equal to the geometric mean of protein abundance. The number of mistranscription errors observed is expected to follow a binomial distribution with *r* trials, each with probability *p* of an error. We thus estimated a standard error of 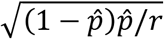, where *r* is the total number of reads within the bin and 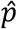 is the observed error frequency within the bin. To generate the 95% confidence interval we multiplied this standard error by 1.96. To keep standard errors for low-abundance bins reasonably low, data from several low-abundance bins were combined.

Binned data is shown for the purpose of illustrating that it is appropriate to log-transform protein abundance before using it as a linear predictor of error rate. Note that it is normal for the edge bins to depart from the linear trend (Wilke 2013), and thus the linearity of the fit should be judged within the central region of the relationship.

## Acknowledgements

This work was supported by the Undergraduate Biology Research Program at the University of Arizona, the John Templeton Foundation [39667, 60814], and the National Institutes of Health (GM104040). We thank Christophe Herman for helpful discussions and Charles Traverse and Jean-François Gout for making their preprocessed data readily available to us.

