## Supplementary Figures 1-3 for "High transcriptional error rates vary as a function of gene expression level"

**Supplementary Figures for**
**"High transcriptional error rates vary as a function of gene expression level"**
**by Meer et al.**

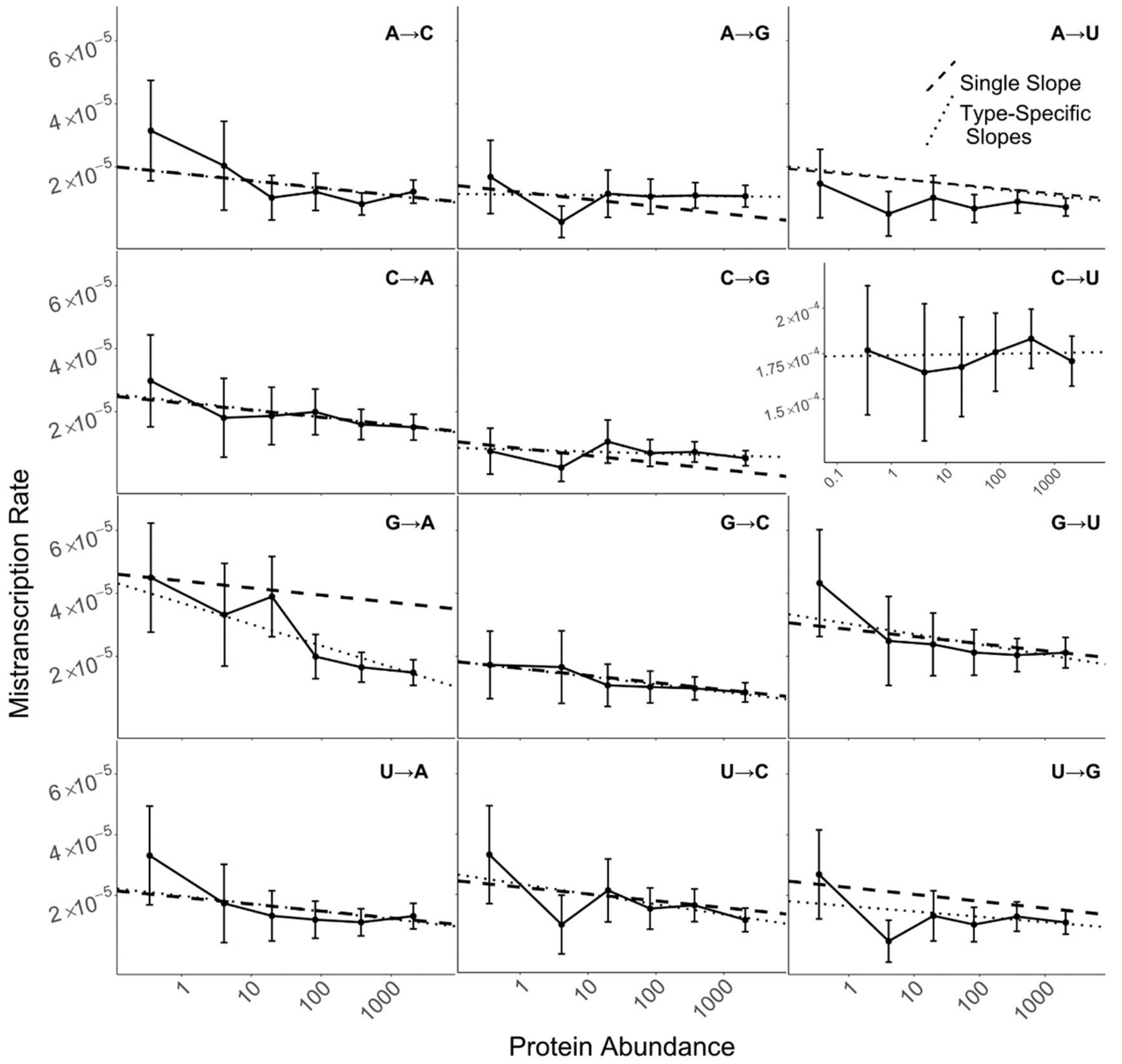


**Figure S1. While different substitution types occur at different rates, all but C→U occur less often for highly abundant *E. coli* proteins.** Dashed lines show our Eq. 1 model, in which the slope is the same across 11 error types (all but C→U). To display this model, we averaged the intercept over the four conditions, weighted according to the numbers of reads in each condition. Dotted lines show linear models with both the slope and intercept fitted separately for each error type using data pooled across all four conditions. Solid lines show the mean mistranscription rates, binned by protein abundance as described in the Methods, plotted according to mean protein abundance within each bin; error bars show 95% CI. Data were divided into 8 bins; because of the limited availability of reads for low-expression genes, data within the first three bins were pooled. Note that mistranscription rate is per possible error, so total mistranscription rate per nucleotide is around three times larger. The C→U error plot is shown inset due to very different y-axis values.

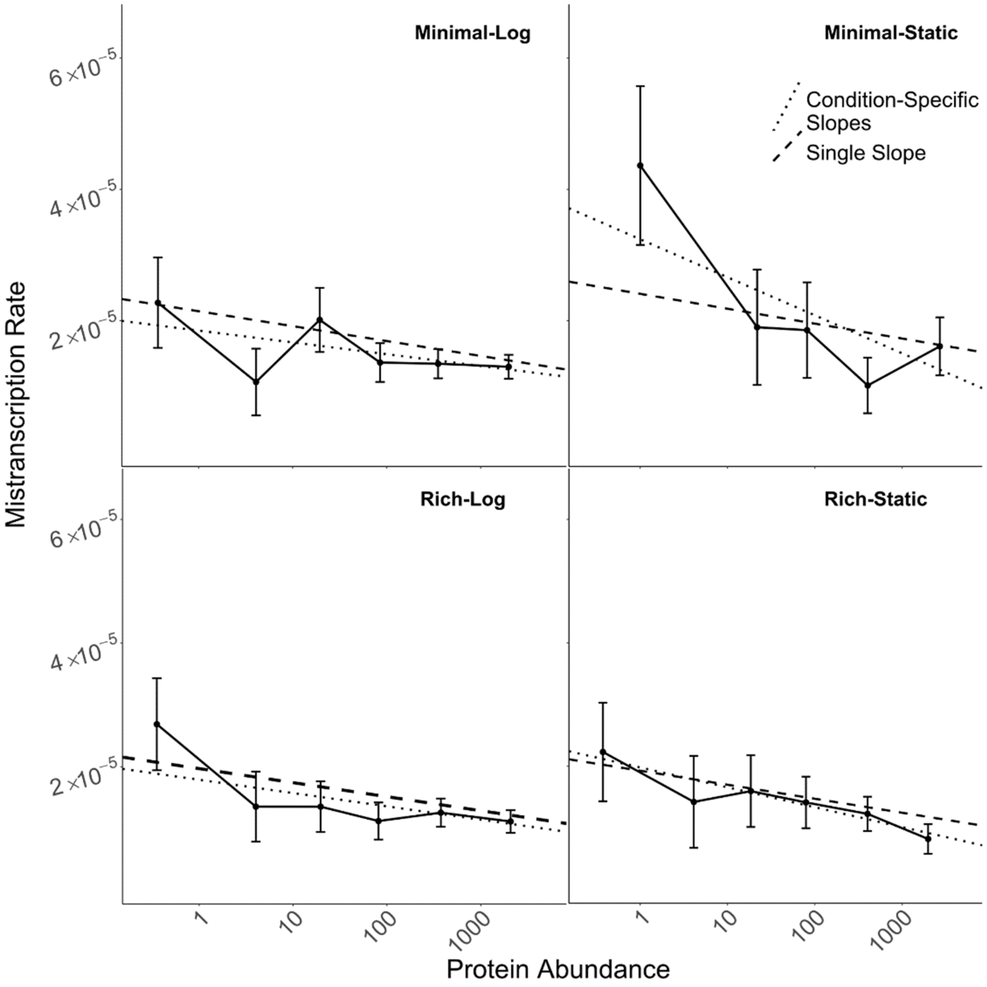


**Figure S2. Different experimental conditions for *E. coli* yield similar mistranscription rates.** Dashed lines show our Eq. 1 model, in which the slope is constant across all conditions. To display this model, we averaged the intercept over all error types, weighted by the frequencies of opportunity for the error type to occur (i.e. by the numbers of reads of sites with A/C/G/U). Dotted lines show linear models with both the slope and intercept fitted separately for each condition using data pooled across all error types except C→U. Solid lines show the mean mistranscription rates, binned by protein abundance as described in the Methods, plotted according to mean protein abundance within each bin; error bars show 95% CI. Data were divided into 8 bins; because of the limited availability of reads for low-expression genes, data within the first three bins were pooled. Note that mistranscription rate is per possible error, so total mistranscription rate per nucleotide is around three times larger.

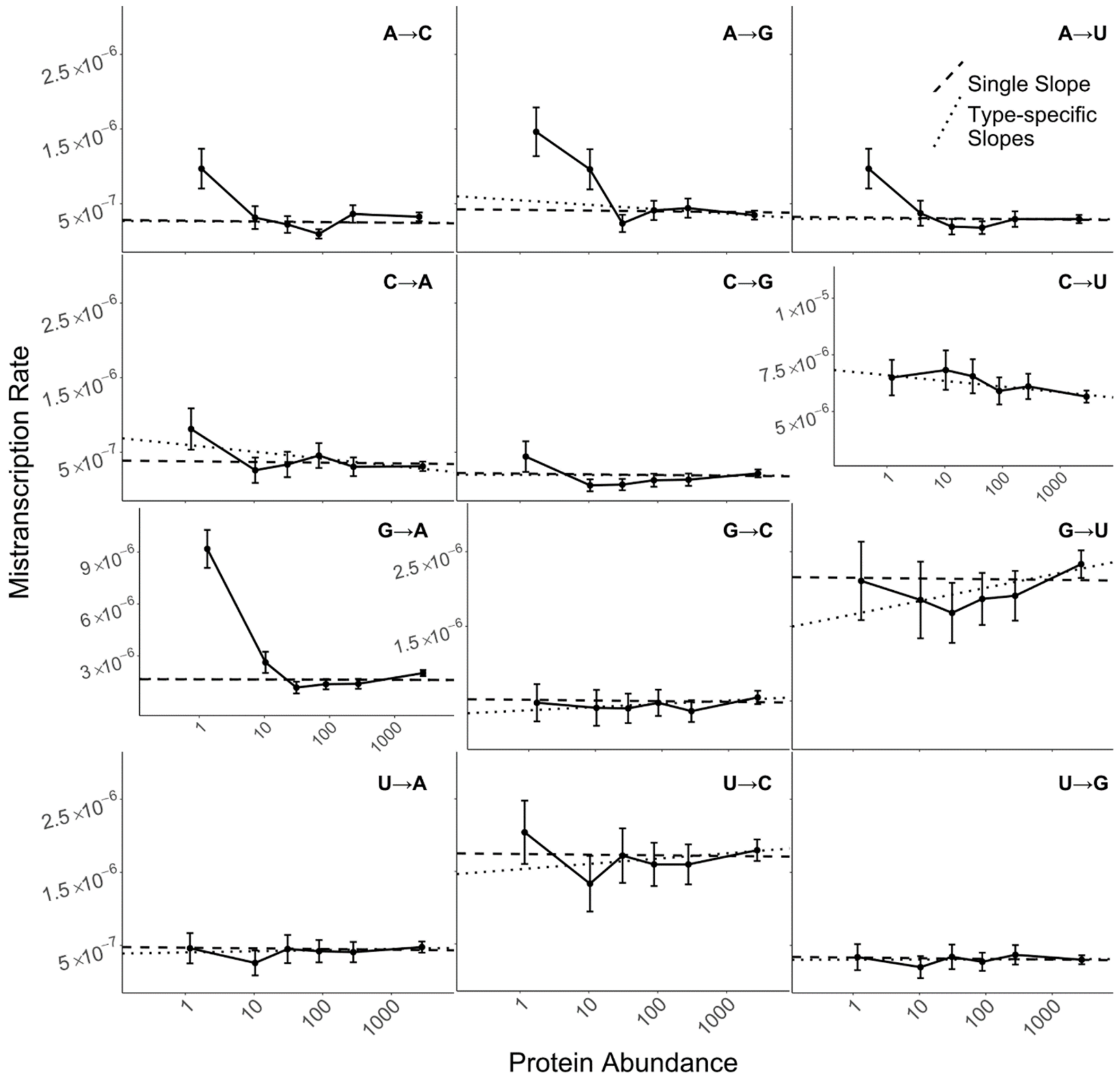


**Figure S3.** ***S. cerevisiae*** **mistranscription rates broken down by base-substitution error types.** Dotted lines show our Eq. 1 model, in which the intercept varies by error type, but the slope is the same across all error types. Dashed lines show linear models with both the slope and intercept fitted separately for each error type. Solid lines show the mean mistranscription rates, binned by protein abundance as described in the Methods, plotted according to mean protein abundance within each bin; error bars show 95% CI. Data were divided into 8 bins; because of the limited availability of reads for low-expression genes, data within the first three bins were pooled. Note that mistranscription rate is per possible error, so total mistranscription rate per nucleotide is around three times larger. The C→U and G→A plots are shown inset due to radically different y-axis values than for the other error types. Despite the G→A outlier, a different slope for G→A is not statistically supported ($p$=0.95), and indeed is superimposable with the overall slope. A model with separate slopes for each error type reveals that A→G and C→A have significantly negative slopes ($p = 5\times10^{-3}$ and $p = 5\times10^{-5}$, respectively). However these slopes are very shallow ($-2.6\times10^{-8}$, $-4.1\times10^{-8}$), corresponding to less than 1 fewer error per 10 million protein products.
